# Chemical signatures of social information in Barbary macaques

**DOI:** 10.1101/2024.09.24.614500

**Authors:** Brigitte M. Weiß, Claudia Birkemeyer, Marlen Kücklich, Anja Widdig

## Abstract

Primates are well-known for their complex social lives and intricate social relationships, which requires them to obtain and update social knowledge about conspecifics. The sense of smell may provide access to social information that is unavailable in other sensory domains or enhance the precision and reliability of other sensory cues. However, the cognition of social information in catarrhine primates has been studied primarily in the visual and auditory domain. We assessed the social information content of body odor in a catarrhine primate, the Barbary macaque (*Macaca sylvanus*), in three semi-free ranging groups at Affenberg Salem, Germany. In particular, we related socially relevant attributes (identity, genetic relatedness, rank, sex, age, group membership) to chemical profiles of body odor. We applied non-invasive active sampling via thermal desorption tubes and analyzed samples by gas chromatography–mass spectrometry. We found robust evidence for individual odor signatures and limited support for kin signatures. Chemical profiles were also related to group membership, but little to rank, sex or age. The detected associations between chemical profiles and socially relevant attributes form the theoretical foundations for olfactory information transfer, highlighting the potential of body odor to provide valuable social information in this highly visually oriented primate.

## Introduction

The development and maintenance of social relationships requires animals to acquire and update social information about conspecifics. For instance, animals that form stable relationships need to be able to recognize the unique features of other individuals, and indeed, individual recognition is widespread in the animal kingdom ^e.g. 1–3^. Additionally, social animals may interact differently with each other depending on attributes such as sex, rank, group membership or genetic relatedness ^4^.

Primates are particularly well-known for forming intricate social relationships and frequently show strong and long-lasting relationships among kin ^5^, but also with unrelated individuals ^6,7^.

Representations of these social relationships are based on the recognition of traits such as identity, rank and kinship ^8^. For instance, rhesus macaques (*Macaca mulatta)* discriminate between individuals, and between familiar as well as unfamiliar kin ^1,9,10^. Various macaque species also appear to classify their group members’ social relationships with others, suggesting that they attend to interactions in which they themselves are not involved (i.e. third-party relationships ^see^ ^11^). Similarly, baboons discriminate between individuals according to their identity and rank, and they also appear to recognize third-party relationships ^8,11,12^.

The social information underlying differentiated interactions can be emitted and received via different sensory modalities ^13,14^. Olfaction is among the evolutionarily oldest of the animal senses ^15^, and social communication is considered as one of its major functions ^16^. Indeed, social interactions are mediated via olfaction in numerous mammals such as African elephants (*Loxodonta africana*) ^17^, black rhinoceros (*Diceros bicornis*) ^18^ and various primates. For instance, many strepsirrhine (lemurs, lorises and galagos) and platyrrhine (South American) primates show pronounced scent-marking behavior ^e.g. 19–21^. Observational, experimental and chemical studies suggest that primate scent marks as well as odors emanating directly from the body may convey a wealth of information about a sender such as identity, sex, age, rank or reproductive state ^e.g. 20,22–24^, and that scent marks primarily serve in intra-sexual competition and mediating reproductive behavior ^13,22,25^.

Notably, vision has gained more and more importance during primate evolution ^see^ ^26^, and catarrhine (African and Asian) primates, in particular, are highly visually oriented animals ^27^. Scent-marking, on the other hand, is uncommon in catarrhines ^28^, and in contrast to evolutionarily older primates, catarrhines lack a functional accessory olfactory system ^29^. Accordingly, catarrhine primates have long been regarded as little predisposed for olfactory communication ^30^. However, the main olfactory system too has an important part in the processing of cues relevant to social recognition, rendering the lack of a functional accessory olfactory system of little significance for the communicative functions of olfaction ^see^ ^16,31^. Furthermore, catarrhine olfactory sensitivity may be on par or even exceed those of other mammals ^32,33^. Research on catarrhine olfactory communication has primarily focused on humans ^34^. Similar to other primates, human body odor has been associated with socially relevant information such as identity, sex, age, kinship or reproductive status (reviewed in ^35^), although the latter is debated ^see^ ^36^. Also in mandrills (*Mandrillus sphinx*), one of the few catarrhine species known to have scent glands, body odor has been related to sex, age, dominance status, group membership and, possibly, individual identity ^37,38^. Results of recent studies further show that the body odor of other catarrhine species too contains information on socially relevant attributes such as sex, rank, female reproductive status, genetic relatedness or group membership may be recognized and acted upon by conspecifics (rhesus macaques, *M. mulatta* ^39,40^ chimpanzees, *Pan troglodytes* ^41^, olive baboons, *Papio anubis* ^42^, Japanese macaques, *M. fuscata* ^43^). Nonetheless, with the exception of a few species such as humans or mandrills, olfactory communication remains severely understudied in catarrhine primates ^30^. This is particularly true for chemical profiles of body odor (i.e. the presence/absence and relative abundance of chemical compounds), which restricts our understanding of which olfactory cues may be available to conspecifics within catarrhines at all. To reduce this gap, we assessed the social information content of (i.e. potential olfactory cues in) body odor in a catarrhine primate by relating chemical profiles of body odor to socially relevant attributes of Barbary macaques living in natural social structures under semi-free conditions at Affenberg Salem, Germany.

Like other macaque species, Barbary macaques live in multi-male, multi-female groups with female philopatry and male dispersal ^44,45^. Matrilineal kin (especially mother-offspring and maternal half-sibling dyads) form strong bonds throughout their lives and are therefore highly familiar with each other ^5,46^. Patrilineal kin (e.g. father-offspring or paternal half-sibling dyads), on the other hand, typically are less closely associated and may even reside in different groups so that they can be expected to be less familiar with each other ^see^ ^47,48^. Females show matrilineal rank acquisition in which female relatives take similar ranks ^49,50^. Adult males form dominance hierarchies in which older males tend to rank higher, but males beyond their prime (i.e. older than 14-16 years) are subordinate to prime age males due to being in inferior physical condition ^51^. Male Barbary macaques also form strong, enduring affiliative relationships often referred to as “friendships” ^52^, whereby triadic interactions between two males and an infant represent an important form of male affiliative interaction ^53^.

Various studies suggest that Barbary macaques can discriminate between individuals, sexes, age classes and group members using visual or acoustic cues ^54–56^. They further distinguish familiar maternal kin from non-kin using acoustic cues ^57^. However, Barbary macaques, which are considered to be mating particularly promiscuously ^58^, do not seem to differentiate between unfamiliar kin and non-kin either in a mating context ^47^ or in infant care ^53,59^. It remains unclear whether this is due to i) a lack of kin-related cues, ii) Barbary macaques not being able to recognize available cues or iii) Barbary macaques recognizing kin-related cues but not acting upon them. Kin-related cues in body odor composition have been shown in a closely related species (rhesus macaques) ^40^, and Barbary macaques also use olfaction in social contexts ^60^, but systematic studies on olfactory kin signatures or other socially relevant information in Barbary macaque body odor are, to date, absent.

In the present study we aimed to i) identify variation in body odor composition (i.e. presence and abundance of chemicals) that is systematically associated with socially relevant information about individuals, and ii) to pinpoint potential semiochemicals (i.e. compounds and compound mixtures involved in chemical interactions between organisms^15^) associated with the respective traits. Our particular focus was on traits that form the basis for differentiated social relationships, i.e. individual identity, genetic relatedness between and social rank of individuals. We further assessed effects of sex, age and group membership on chemical profiles of body odors while statistically controlling for potential seasonal and methodological variation (such as sampling distance and volume, or analytical batch) in the data. We hypothesized that the odorants emanating from the skin and fur of Barbary macaques contain socially relevant information. In particular, we expected the similarity between chemical profiles to be greater within than between individuals, and higher among related than among unrelated individuals. Because maternal kin are in frequent physical contact and may exchange odors from the environment or share more of the odor-producing microbiome, we expected maternal kin to exhibit stronger kin signatures than paternal kin, for whom kin signatures should primarily result from similarities in their genetic architecture. We also expected individuals of similar rank to have more similar chemical profiles and/or to show rank-related differences in the relative abundance of particular compounds, as also suggested in ^61^, because individuals of similar rank may have similar hormonal profiles (e.g. androgens or glucocorticoids), access to food resources and/or space use ^see^ ^62–65^. Finally, given the comprehensive evidence for systematic differences in body odor composition with respect to sex, age and group membership in other primates, we also expected similarities between chemical profiles and/or the relative abundance of compounds to be associated with these traits.

## Methods

### Study animals

The study was conducted at Affenberg Salem, Germany, home to ∼ 200 Barbary macaques in three social groups roaming year-round within a 20 ha forested enclosure. The monkeys are well-habituated to humans, as the park is open to visitors from spring to autumn, and to researchers year-round (see ^66^ for details on the park). The monkeys’ diet consists of plants and insects found naturally within the park as well as fruit, vegetables and grain that is provided daily by park staff. Ponds and water troughs are distributed throughout the park and grant *ad libitum* access to water. Individuals differ in facial pigmentation patterns, coat color and stature and have an alphanumeric code tattooed on the inner thigh, which makes them well distinguishable for experienced observers. Park staff monitor the groups on a near-daily basis and record major life history events such as births, deaths or group changes. About two thirds of the adult females receive hormonal implants (Implanon NXT) to control population size.

We collected odor samples from a total of 72 focal males and females from all three social groups (“C”, “F” & “H”) and across all ages (2 - 27 years, mean ± SD = 10.5 ± 5.4 years) that were most suitable to address the main study questions. Focal animals included 13 females without and 18 females with hormonal contraception throughout the study period; two females received contraception for part of the study period (see Supplement 1 Tab. S2 and statistical analyses section below). To assess individual odor signatures, we included 24 particularly well-habituated reproductively mature individuals (> 4 years) that could be sampled frequently (10 samples per ID, Pollard et al. 2010). To address kinship, we included mothers/fathers and their offspring (> 2 years, because younger individuals could not be reliably habituated to sampling yet) from a total of 12 maternal and 12 paternal kin units. For rank of males and females, we selected individuals from the highest- and lowest-ranking adults in their respective groups (17 individuals per rank category, see details in statistical analysis section).

### Odor sampling

One experimenter (B.M.W.) collected 476 odor samples from the focal animals (mean ± SD = 6.6 ± 2.6, median = 5, range 1 – 11 samples per ID) in five sampling periods between October 2020 and April 2022. Sampling took place in spring (N = 213 samples from 51 individuals), summer (N = 113 samples from 39 individuals) and autumn (N = 150 samples from 56 individuals), but, for logistic reasons, not during the peak of the mating season in the months of November, December and January. As Barbary macaques, like other macaque species, do not appear to have specialized scent glands, we non-invasively collected the airborne volatile and semi-volatile compounds emanating from the skin or fur of the animals across different parts of the body (see below). We employed active sampling with thermal desorption (TD) tubes (stainless steel, Supelco), which contained two adsorbent polymers (Tenax TA and XAD-2, Sigma Aldrich) that trap a wide range of chemical compounds ^67^. The tubes were connected to a silently operating air pump (BiVOC2, Holbach) via a silicone hose ^68^. To be able to trap the air in the immediate vicinity of the animals (1 to 15 cm, mean ± SD = 3.5 ± 2.6 cm), monkeys were habituated to retrieve pieces of peanuts from a pipe collar (7.5 cm diameter) strapped to a horizontal branch or rail while the experimenter stood next to the setup and pointed a wooden stick with the TD tube mounted at its end towards the monkey. Once a monkey calmly started to retrieve the peanuts, the experimenter opened the TD tube, positioned it close to the monkey and started the air flow through the tube with the flow rate set to 1 L/min.

Sampling was resumed until an air volume of 0.5 – 1 L was collected. If the animal moved away before the sample was completed, the pump was immediately stopped, the tube closed, the volume already sampled noted and sampling resumed as soon as possible onto the same TD tube (mean ± SD = 2.3 ± 1.5 subsamples). A sample was typically completed with < 5 min between subsamples, except for 10 of the 476 samples in which the sample was completed on different days (see Supplement GCMSdata). Samples were collected from body parts that could be reliably sampled without disturbing the animals, which primarily constituted the shoulders, flanks or outer thighs (54 % of samples being collected exclusively from these regions, and 86 % of samples included these regions). Whenever possible we avoided sampling the hands, feet or facial area, which were the body parts most likely contaminated with food, feces or other items from the environment. To be able to statistically control for potential variation arising from the sampled body parts we grouped them into 4 categories: “upper” constituted samples collected from the upper part of the body (i.e. arms, shoulders and/or the back of the head), “central” the middle part of the body (i.e. flanks, central back or chest), and “lower” the lower part of the body (i.e. hips, thighs and legs). Finally, we grouped all samples collected from multiple regions of the body (i.e. upper, central and/or lower) into the category “mixed” (see Supplement GCMS data).

We avoided sampling when conspecifics were within 3 m of the target animal or higher-ranking individuals were within sight, when animals had body contact with group members less than 5 minutes before ^68^, when the monkeys’ fur was moist or wet, or when it was very windy. To avoid disturbances during sampling and contamination with human odors, we did not collect any samples when visitors or park staff were within 15 m. All equipment was handled wearing disposable gloves. The experimenter wore a medical mouth-nose cover during sampling and avoided the use of perfumed products or strongly smelling foods before and on sampling days.

To control for chemical compounds originating from the sampling material and sampling environment *per se*, we further collected samples of air in the absence of animals (N = 17), TD tubes mounted for sampling in the absence of animals but without drawing air through the tube (N = 6) and TD tubes taken to the field but remaining closed at all times (N = 9) as blanks. Before each use, we cleaned the TD tubes in a thermal conditioner (TD Clean-Cube, Scientific Instruments Manufacturer) under a constant stream of nitrogen for 120 minutes. When not in use, the tubes were closed tightly with Swagelok brass caps, wrapped in aluminum foil and stored in airtight bags at ambient temperature.

### Chemical analysis & profiling

Samples were analyzed with gas chromatography – mass spectrometry (GC-MS) on a Shimadzu TQ8040 GC-MS coupled to a thermal desorption unit (TD-20) with a scan range from 30 to 500 *m/z* following established GC-MS protocols for the analysis of TD tube samples ^67,68^. In the process, the trapped organic compounds were entirely desorbed from the tube, which then was conditioned and re-used for sampling. To remove any potential residues from previous measurements from the column, we ran three blank samples at the start and end of each batch as well as after every tenth sample.

GC-MS data were processed following an established, semi-automated procedure ^67,68^, in which a preliminary compound library was created through automated peak detection and alignment of peaks by retention time using AMDIS (version 2.71 ^69^). We manually corrected this library by taking into account characteristic mass-to-charge ratios of peaks in addition to retention times. We subsequently characterized all unique compounds by their retention time and mass spectral patterns but did not attempt to chemically identify them unless their abundance was later determined to be statistically associated with particular traits (see below). For the 264 compounds in this library, we extracted the peak areas for the highest ion not present in adjacent peaks using the Shimadzu GCMS Browser software. In a subsequent data cleaning step we excluded 81 compounds consistently found in blanks with peak areas higher than in the animal samples from further analysis, as they are unlikely to have derived from the target animal ^detailed^ ^in^ ^68^, and performed statistical analyses using the remaining 182 compounds (see Supplement 2 for full list of retention times and characteristic mass-to-charge ratios). Furthermore, we assigned undetected peaks, peaks with a signal to noise ratio < 1 and with a peak area below the noise threshold (visually determined to be at 1000) a peak area of 0, because these peaks were not reliably separable from background noise. All these steps were done based on uninformative sample numbers and thus blind to the social information about the sampled individuals. We tentatively identified compounds statistically related to particular traits (see below) by comparing their mass spectra with the best matches of the NIST Mass Spectral Library (NIST14; National Institute of Standards and Technologies, Gaithersburg, MD, USA).

### Parentage analysis

Pregnancies and births have been monitored consistently by Affenberg staff since the 1970s, so that partial pedigrees could be built tracing back maternal relatedness over 5-6 generations for most individuals based on these long-term observational records.

For genetic parentage analysis we used DNA extracted from blood, tissue, hair or feces of 144 individuals (including 68 of the 72 odor-sampled animals, see Supplement 1 for details on genetic samples and paternity analysis). We genotyped all individuals at an average of 16.7 ± 2.8 microsatellite loci using a marker panel previously established for the study population ^70^ (see Supplement 1, Tab. S1).

By applying a combination of exclusion and likelihood methods we were able to assign fathers to 23 of the 28 offspring from cohorts 2017-2019 and to 18 individuals initially sampled as mothers or potential sires in the present study and born between 2011 and 2016 (see Supplement 1 for details). For the 68 odor-sampled and genotyped individuals, we were able to genetically confirm 25 fathers and 41 mothers.

Based on the genetic analysis and maternal pedigree data from long-term records, we calculated pedigree relatedness (r) per kin line (maternal or paternal/mixed) for each possible dyad in our study using the software TRACE (version 0.1.0) ^71^. Maternal r reflected genetic relatedness between two individuals of either sex exclusively via female ancestors, while paternal/mixed r (hereafter just called “paternal”) reflected any connections involving at least one male ancestor linking two given individuals. Because no genetic samples could be obtained for potential sires deceased before the onset of the study, paternities could only be reliably determined for the younger cohorts, and therefore our values of pedigree relatedness for the paternal line are likely underestimated in this analysis.

### Dominance and rank

At the onset of the study, we asked experienced park staff to classify individuals from low to high-ranking, and to identify the alpha male and female in each group based on observed agonistic interactions and priority of access to resources. This initial judgement was used to pre-select focal individuals for habituation to the sampling setup. We further conducted dynamic Elo-rating ^72^ using *ad libitum* observations of agonistic interactions (as defined in ^73^) throughout all sampling periods.

Dynamic Elo-rating allows for incorporating intensities of interactions as well as prior knowledge and therefore requires fewer observed interactions and interacting dyads than unmodified Elo-rating ^74^ or matrix methods to obtain robust rank estimates ^72^. We used the initial classification by park staff, supplemented by a classification by B.M.W., to define starting values for Elo-rating, and updated this information with 462 female-female and 240 male-male interactions of varying intensities observed between October 2020 and April 2022. We computed ranks separately per sex and group, using the observations during the first two sampling periods as a “burn-in” phase, and extracted the ranks of the focal individuals at the ends of the second to fifth sampling period for further analysis.

### Ethical note

All experimental protocols were approved by Affenberg Salem and complied with the ARRIVE guidelines. All samples collected specifically for this study (i.e. odor and fecal samples) were collected non-invasively and the animals participated in the odor sampling procedure voluntarily. In line with the animal protection law in Germany, we therefore required no further animal ethics approval. All methods were performed in accordance with the relevant guidelines and regulations.

### Statistical analysis

#### General procedures

We conducted all analyses in R (version 4.3.3) ^75^.

To investigate chemical similarity between samples, we performed permutational multivariate analyses of variance (perMANOVAs) using the function “adonis2” from the package “vegan” (version 2.6-4)^76^. These analyses relate pairwise chemical (dis)similarities between all samples entered into a given analysis to (dis)similarities between traits using distance matrices automatically assembled from the sample-specific information provided. As measure of distance (or dissimilarity) we used Bray-Curtis indices calculated from standardized, log(x+1)-transformed peak areas ^68,77^. Depending on the investigated trait(s) we used different data subsets and predictor variables as described for the different traits below. Unless noted otherwise, we set ID as stratum to account for repeated measures per individual. For perMANOVAs with multiple test predictors we first conducted a full-null model comparison by setting the argument “by” to “NULL”, and only if this was significant (p < 0.05) we investigated the effects of the single predictor variables by setting the argument “by” to “margin”. In case of non-significant interactions, we re-ran the marginal model without the interaction to be able to assess the effect of the respective main terms. We ran all perMANOVAs with the default of 999 iterations.

When the analysis of chemical (dis)similarities involved predictor variables describing pairs of samples (e.g. pedigree relatedness) and/or required the selection of specific pairs of samples (which is not possible with the perMANOVA), we further conducted multi-membership models. Multi-membership models extend multilevel models by allowing to fit data with multiple membership structures that are not purely hierarchical ^78^, such as terms whose levels are present in more than one variable (e.g. ID in either sample 1 or sample 2 of a given pair-wise comparison). For these models we computed Bray-Curtis dissimilarities from standardized, log(x+1)-transformed peak areas using the package “vegan” and, for easier interpretation, transformed them into similarities by subtracting the dissimilarity scores from 1. We fitted these models with chemical similarity as response variable and various test and control predictors as described below. As similarity scores follow a beta distribution, we computed the models using a Bayesian approach in the package “brms” (version 2.21.0) ^79^, which is the only package we are aware of that implements multi-membership models with beta distribution. We used the default settings for priors, i.e. uninformative priors.

We further conducted compositional mixed model analyses, suitable for data whose components are constrained to sum to a constant value, for a detailed analysis of chemical profiles that considers the peak areas of all compounds per sample rather than overall (dis)similarities between samples, and may therefore aid in detecting (suites of) compounds associated with particular traits. For this purpose, we vectorized the data matrix and included the matrix rows and columns (i.e. the samples and compounds) as random intercepts to avoid pseudo-replication and heteroscedastic variance ^68,80^. As response variable we used standardized peak areas. Because standardized areas are bound between 0 and 1 and sum up to 1 per sample, we applied a centered log-ratio (clr) transformation ^81^ and modelled the transformed values with a Gaussian error distribution. As effects of interest in this analysis we modelled the interactions between the predictors and compound, i.e. the random slopes & random interactions of fixed effects test predictors within compound ^see^ ^68^. To determine which compounds were associated with particular traits, we subsequently extracted the compound-specific random slopes and random interaction estimates for each trait. We considered a given compound to be robustly associated with a trait if its credible intervals did not overlap zero (in case of continuous predictors) or did not overlap between trait levels (in case of categorical predictors). To follow a consistent modelling approach, we also performed this model in the package brms.

In all mixed models we z-transformed continuous predictor variables to a mean of 0 and an SD of 1 prior to fitting the model to facilitate convergence and interpretation of model estimates ^82^. We assessed collinearity between predictor variables using the function “vif” from the R package “car” ^version^ ^3.1–2^ ^83^, whereby we considered values < 4 as unproblematic. To assess model convergence we inspected rhat values and effective sample sizes (ESS), with rhat values deviating from 1 by less than 0.01and ESS values > 400 indicating satisfactory convergence ^84^.

We present model results most relevant for the interpretation of the test predictors in the main text, and full model results, including rhat and ESS values, in Supplement 1 (Tables S3-S6).

We could not account for contraception statistically in any of the analyses given that contraceptive state applied only to a minority of samples and individuals.

#### Individual identity

To investigate individual signatures, we analyzed distance matrices using a perMANOVA with ID as test predictor while controlling for sex, age (in years), group and season of the year (spring, summer or autumn). Because ID was the test predictor, we did not set ID as stratum. For this analysis we used data from 24 individuals (12 males, 12 females) sampled 10 times each. As individual signatures commonly result from the unique combination and abundance of compounds (i.e. the overall similarity between chemical profiles) rather than certain compounds that vary consistently between all individuals ^85,but^ ^see^ ^43^, we did not conduct a more detailed analysis of chemical profiles (compositional models) for individual identity and consequently also did not attempt to identify specific compounds associated with individual odor signatures.

#### Genetic relatedness

Similar to ID, kin signatures should result from similarities in the overall composition of compounds rather than specific compounds. We therefore assessed whether the chemical (dis)similarities between samples are explained by genetic relatedness. In the first of two sets of analyses, we computed perMANOVAs using only samples of genetically confirmed units of close kin. Each kin unit comprised 2 – 4 individuals in up to 3 generations and as such, parent-offspring, grandparent-grandchild and/or half-sibling dyads, reflecting close kin with relatedness coefficients ranging from 0.25 to 0.5. We ran the perMANOVAs separately per kin line, using the maternal or paternal kin unit (i.e. belonging to the same or a different kin unit) as test predictor, while controlling for sex, age and season of the year (spring, summer or autumn). All individuals used in these analyses were sampled for odor at least 5 times. For maternal kin we used 220 samples from 31 individuals and for paternal kin 179 samples from 29 individuals in 12 kin units per kin line (see Supplement 1, Tab. S2). Maternal/paternal relatedness between individuals from different units ranged from 0 to 0.125, reflecting distant or non-kin.

While belonging to a certain kin unit was genetically confirmed in this first set of analyses, it treated relatedness categorically as belonging to a given kin unit or not, but lacked the possibility to differentiate the pair-wise degree of relatedness. In a second analytical approach we therefore conducted a multi-membership model to relate pair-wise chemical similarities between samples to kinship while accounting for a variety of fixed and random predictor variables. For the multi-membership model we used the data from all genotyped and odor-sampled individuals (N = 68), irrespective of whether their maternities or paternities were genetically solved. We fitted dyadic Bray-Curtis similarities as beta-distributed response variable and relatedness between members of a dyad as two separate test predictors: one reflecting maternal kinship (i.e. individuals linked exclusively via the maternal line) and one reflecting paternal & mixed kinship (i.e. containing at least one male linking two individuals, which breaks the familiarity between maternal kin typical for matrilineally structured primate groups). To control for other sources of biological variation we also fitted the age and rank difference between the two individuals in a given dyad, the number of days between the two samples and the difference in the time of day the sample was collected. We further included whether or not the two individuals were of the same sex, belonged to the same group, were sampled in the same season of the year and from the same body region. To control for variation in sampling and measurement we included the difference in sampling distance, sampling volume, the number of subsamples needed to complete a sample and whether or not samples were analyzed in the same GC-MS batch. To account for repeated measures we fitted the sample, the individual and the dyad (i.e. the unique combination of individuals) and random intercepts, whereby sample and individual were fitted as multi-membership terms because a given sample or individual could occur in either of the two columns describing the elements of a dyad. For a more accurate estimation of fixed effects, we further included all technically possible random slopes of fixed effects within the random intercepts of individual and dyad. Using 9000 post-warmup draws, the model converged with all rhat values deviating from 1 by < 0.01 and all ESS > 400 (see Table 1).

**Table 1:**
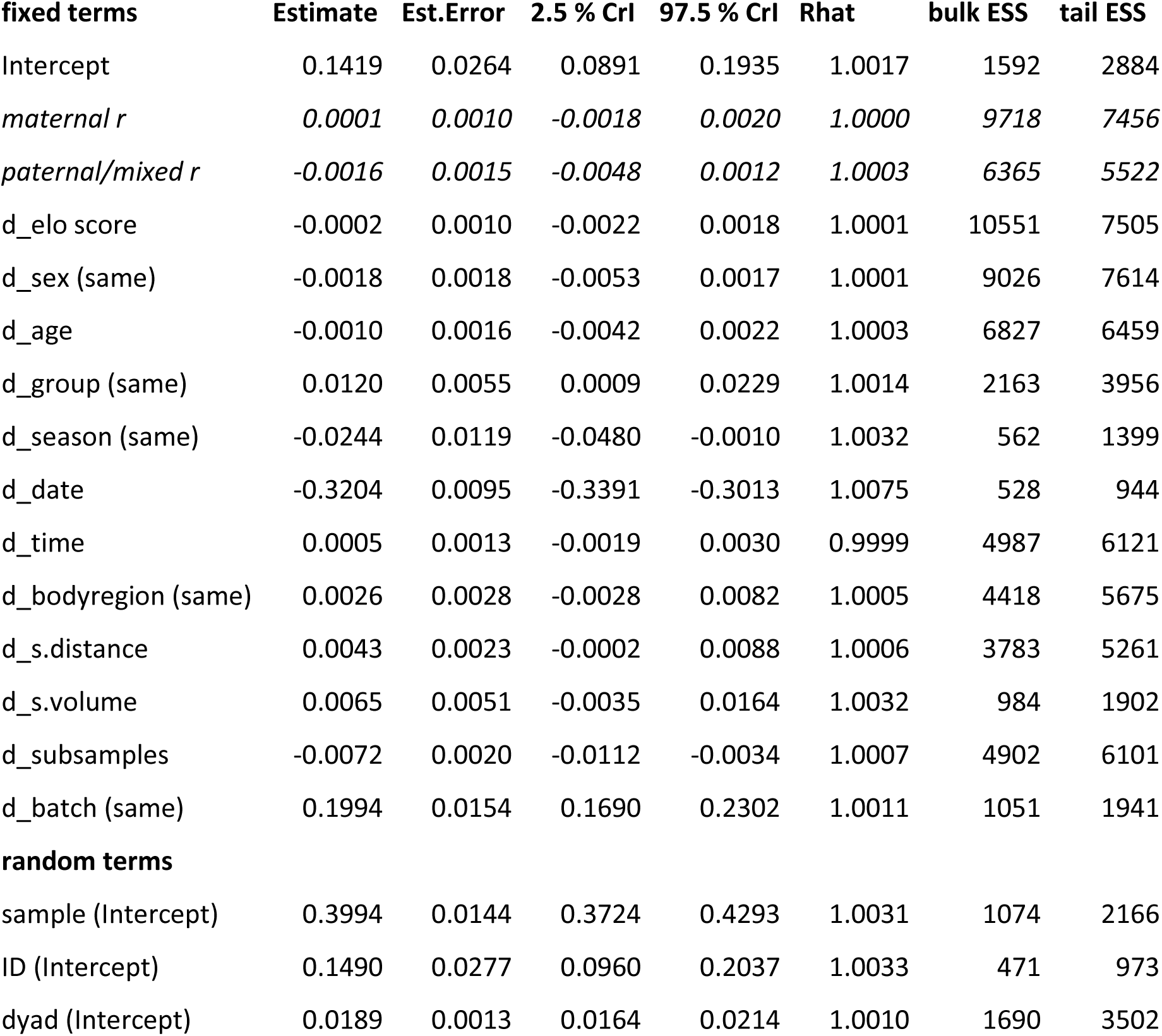
Estimates for fixed effects and random intercepts from the multi-membership model on genetic relatedness. Terms of primary interest are indicated in italics. CrI = Credible Interval. ESS = effective sample size.

| **fixed terms** | **Estimate** | **Est.Error** | **2.5 % CrI** | **97.5 % CrI** | **Rhat** | **bulk ESS** | **tail ESS** |
| --- | --- | --- | --- | --- | --- | --- | --- |
| Intercept | 0.1419 | 0.0264 | 0.0891 | 0.1935 | 1.0017 | 1592 | 2884 |
| *maternal $r$* | *0.0001* | *0.0010* | *-0.0018* | *0.0020* | *1.0000* | *9718* | *7456* |
| *paternal/mixed $r$* | *-0.0016* | *0.0015* | *-0.0048* | *0.0012* | *1.0003* | *6365* | *5522* |
| d_elo score | -0.0002 | 0.0010 | -0.0022 | 0.0018 | 1.0001 | 10551 | 7505 |
| d_sex (same) | -0.0018 | 0.0018 | -0.0053 | 0.0017 | 1.0001 | 9026 | 7614 |
| d_age | -0.0010 | 0.0016 | -0.0042 | 0.0022 | 1.0003 | 6827 | 6459 |
| d_group (same) | 0.0120 | 0.0055 | 0.0009 | 0.0229 | 1.0014 | 2163 | 3956 |
| d_season (same) | -0.0244 | 0.0119 | -0.0480 | -0.0010 | 1.0032 | 562 | 1399 |
| d_date | -0.3204 | 0.0095 | -0.3391 | -0.3013 | 1.0075 | 528 | 944 |
| d_time | 0.0005 | 0.0013 | -0.0019 | 0.0030 | 0.9999 | 4987 | 6121 |
| d_bodyregion (same) | 0.0026 | 0.0028 | -0.0028 | 0.0082 | 1.0005 | 4418 | 5675 |
| d_s.distance | 0.0043 | 0.0023 | -0.0002 | 0.0088 | 1.0006 | 3783 | 5261 |
| d_s.volume | 0.0065 | 0.0051 | -0.0035 | 0.0164 | 1.0032 | 984 | 1902 |
| d_subsamples | -0.0072 | 0.0020 | -0.0112 | -0.0034 | 1.0007 | 4902 | 6101 |
| d_batch (same) | 0.1994 | 0.0154 | 0.1690 | 0.2302 | 1.0011 | 1051 | 1941 |
| **random terms** |  |  |  |  |  |  |  |
| sample (Intercept) | 0.3994 | 0.0144 | 0.3724 | 0.4293 | 1.0031 | 1074 | 2166 |
| ID (Intercept) | 0.1490 | 0.0277 | 0.0960 | 0.2037 | 1.0033 | 471 | 973 |
| dyad (Intercept) | 0.0189 | 0.0013 | 0.0164 | 0.0214 | 1.0010 | 1690 | 3502 |

Similar to individual identity we did not conduct compositional models for genetic relatedness and did not attempt to identify specific compounds associated with kin odor signatures.

#### Dominance and rank

To address potential dominance and rank-related signatures, we first conducted a perMANOVA with rank as a categorical test predictor (high/low) in interaction with sex while controlling for age, group and season of the year. For this analysis we used 260 samples from 34 individuals sampled at least 5 times each and classified as particularly high- or low-ranking in their respective groups based on judgements of staff and confirmed with behavioral data (high-ranking: 9 females & 8 males, low-ranking: 8 females & 9 males). To keep confounds between dominance class and kinship arising from matrilineal rank acquisition to a minimum, we chose no more than two females from the same matriline, so that the 17 females used for this analysis represented 12 different matrilines.

We further conducted a multi-membership model using chemical data and rank as a continuous measure (i.e. Elo scores) using 476 samples from all 72 sampled individuals. We only considered same-sex dyads for this analysis, because Barbary macaque males outrank females and Elo scores were thus computed separately per sex. We fitted dyadic Bray-Curtis similarities as beta-distributed response variable and the difference in Elo scores and its interaction with sex as test predictors.

Similar to the model on genetic relatedness we also controlled for potential sources of biological and technical variance by fitting dyad-specific overall relatedness and differences in age, sampling day, time of day, group, season of the year, body region, sampling distance, sampling volume, number of subsamples and GC-MS batch. As above we fitted sample and individual as multi-membership terms as well as dyad as random intercepts, and all technically possible random slopes within individual and dyad. We subsequently removed the interaction between the difference in Elo scores and sex because it explained no variance (estimate 0.0004 +-0.0028, lower/upper CI -0.005 – 0.006) and fitted both as main terms instead. Using 12000 post-warmup draws, the model without interaction converged with all rhat values deviating from 1 by < 0.01 and all ESS > 400 (see Table 2).

**Table 2:** Estimates for fixed effects and random intercepts from the multi-membership model on rank. Terms of primary interest are indicated in italics. CrI = Credible Interval. ESS = effective sample size.

| **fixed term** | **Estimate** | **Est.Error** | **2.5 % CrI** | **97.5% CrI** | **Rhat** | **bulk ESS** | **tail ESS** |
| --- | --- | --- | --- | --- | --- | --- | --- |
| Intercept | 0.1544 | 0.0394 | 0.0756 | 0.2315 | 1.0010 | 2680 | 5074 |
| *d_Elo score* | *-0.0001* | *0.0014* | *-0.0027* | *0.0026* | *1.0002* | *19102* | *9841* |
| total r | -0.0021 | 0.0014 | -0.0049 | 0.0007 | 1.0002 | 18584 | 8977 |
| sex (male) | -0.0228 | 0.0526 | -0.1265 | 0.0798 | 1.0014 | 2879 | 5192 |
| d_age | 0.0013 | 0.0021 | -0.0030 | 0.0054 | 1.0001 | 14446 | 9510 |
| d_group (same) | 0.0159 | 0.0070 | 0.0022 | 0.0297 | 1.0001 | 5497 | 7723 |
| d_season (same) | -0.0267 | 0.0108 | -0.0484 | -0.0059 | 1.0015 | 2736 | 4694 |
| d_date | -0.3250 | 0.0104 | -0.3456 | -0.3053 | 1.0023 | 1088 | 2508 |
| d_time | 0.0005 | 0.0012 | -0.0019 | 0.0030 | 1.0002 | 15821 | 9039 |
| d_bodyregion (same) | 0.0041 | 0.0033 | -0.0024 | 0.0107 | 1.0002 | 11941 | 8932 |
| d_s.distance | 0.0054 | 0.0025 | 0.0006 | 0.0103 | 1.0007 | 11074 | 9460 |
| d_s.volume | 0.0081 | 0.0054 | -0.0026 | 0.0187 | 1.0014 | 2684 | 3564 |
| d_subsamples | -0.0070 | 0.0021 | -0.0113 | -0.0029 | 0.9999 | 15094 | 10231 |
| d_batch (same) | 0.1838 | 0.0182 | 0.1477 | 0.2188 | 1.0011 | 2652 | 5513 |
| **random term** |  |  |  |  |  |  |  |
| sample (Intercept) | 0.3952 | 0.0141 | 0.3684 | 0.4239 | 1.0016 | 2096 | 4665 |
| ID (Intercept) | 0.1509 | 0.0282 | 0.0988 | 0.2076 | 1.0058 | 891 | 1598 |
| dyad (Intercept) | 0.0189 | 0.0019 | 0.0152 | 0.0226 | 1.0016 | 2254 | 4976 |

To consider the possibility that rank information could be encoded by a few specific compounds (rather than or additional to overall similarities), we further included rank as predictor in a compositional mixed model as described in the analyses of sex, age and group below.

#### Sex, age and group

To address potential signatures of sex, age or group membership we conducted a perMANOVA with these terms as test predictors while controlling for season of the year. For this analysis we used all 476 samples from 72 individuals.

To assess to what extent specific (suites of) compounds may encode information about individual traits, we further computed a compositional mixed model. As fixed effects predictors we fitted sex, age, group and rank (Elo scores) along with season of the year, the sampled body part, time of day, sampling distance, sampling volume and the number of subsamples. As random terms we fitted the intercepts of sample, compound, individual, date, GC-MS batch and TD tube used for sampling, as well as all technically possible random slopes, but no random correlations to avoid an overly complex model. Using 8000 post-warmup draws, the model converged with all rhat values deviating from 1 by < 0.01 and all ESS > 400 (see Table 3).

**Table 3:** Results of the perMANOVA investigating effects of sex, age and group on chemical similarities.

| Term | Df | SumOfSqs | R<sup>2</sup> | F | P |
| --- | --- | --- | --- | --- | --- |
| sex | 1 | 0.466 | 0.004 | 2.332 | 0.001 |
| age | 1 | 0.425 | 0.004 | 2.124 | 0.001 |
| group | 2 | 6.106 | 0.051 | 15.273 | 0.001 |
| season | 2 | 17.225 | 0.145 | 43.083 | 0.001 |
| Residual | 469 | 93.756 | 0.787 |  |  |
| Total | 475 | 119.117 | 1 |  |  |

## Results

### Individual identity

Odor samples from the same individuals were significantly more similar than samples from different individuals, with ID explaining ∼ 20 % of the variance in Bray-Curtis indices (perMANOVA, r^2^ = 0.203, df = 21, F = 3.838, p = 0.001, example chromatograms in Supplement 1, Fig. S1). An individual chemical signature was further supported by the posterior mean estimates for ID in the multi-membership models on genetic relatedness and rank (see Tables 1 and 2).

### Genetic relatedness

Odor samples from close maternal kin were significantly more similar than samples from different kin units (perMANOVA, r^2^ = 0.099, df = 11, F = 2.537, p = 0.001). Furthermore, individuals from the same paternal kin unit tended to have more similar chemical profiles than individuals from different kin units (perMANOVA, r^2^ = 0.132, df = 11, F = 2.772, p = 0.067).

When relatedness was treated as a continuous variable (i.e. using pedigree r) across all genotyped individuals in the data set, neither maternal nor paternal relatedness robustly affected similarities in chemical profiles (Table 1).

### Dominance and rank

The interaction between dominance classes (high vs. low) and sex (male vs. female) had no significant effect on the similarity between chemical profiles (perMANOVA, r^2^ = 0.003, df= 1, F = 0.932, p = 0.982), but individuals from the same dominance class had slightly more similar chemical profiles than individuals from different dominance classes (perMANOVA, r^2^ = 0.002, df = 1, F = 0.879, p = 0.008). When treated as a continuous variable across all individuals in the data set, rank (Elo score) had no robust effect on similarities between chemical profiles (Table 2).

Also the specific composition of chemical profiles showed no pronounced variation in relation to rank (see random slope of Elo score within compound in Table 4), and none of the compounds was robustly associated with rank (see random slopes estimates in Supplement 3).

**Table 4:** Estimates for random slope terms within the random intercept of compound for the compositional model. Terms of primary interest are indicated in italics. CrI = Credible Interval. ESS = effective sample size.

| **slope in compound** | **Estimate** | **Est.Error** | **2.5 % CrI** | **97.5% CrI** | **Rhat** | **bulk ESS** | **tail ESS** |
| --- | --- | --- | --- | --- | --- | --- | --- |
| *Elo score* | *0.0166* | *0.0118* | *0.0007* | *0.0435* | *1.0020* | *3304* | *3855* |
| *sex* | *0.1023* | *0.0191* | *0.0626* | *0.1380* | *1.0006* | *2819* | *3082* |
| *age* | *0.0524* | *0.0175* | *0.0121* | *0.0831* | *1.0017* | *1709* | *1218* |
| *group* | *0.2901* | *0.0157* | *0.2606* | *0.3214* | *1.0012* | *3772* | *4889* |
| season | 1.0605 | 0.0417 | 0.9822 | 1.1461 | 1.0019 | 2698 | 4371 |
| time | 0.0397 | 0.0182 | 0.0041 | 0.0724 | 1.0032 | 1657 | 2624 |
| bodyregion | 0.1333 | 0.0176 | 0.0987 | 0.1672 | 1.0009 | 2690 | 3865 |
| s.distance | 0.2624 | 0.0177 | 0.2290 | 0.2983 | 1.0003 | 2989 | 5028 |
| s.volume | 0.8104 | 0.0420 | 0.7341 | 0.8990 | 1.0015 | 1140 | 2388 |
| subsamples | 0.1280 | 0.0147 | 0.0993 | 0.1569 | 0.9999 | 3441 | 5449 |

### Sex, age and group

When testing sex, age and group membership, the set of predictor variables significantly affected chemical similarities (perMANOVA, r^2^ = 0.213, df = 6, F = 21.145, p = 0.001). In particular, sex, age as well as group explained significant, albeit small proportions of the variance in Bray-Curtis indices (3.5 – 5 %, Table 3). In comparison, season of the year (included as a control predictor) explained more variance than sex, age and group together (14.5 %, Table 3).

This pattern was similar when considering chemical profiles in detail. Slope estimates from the compositional model suggest that the effects of sex, age and group on relative peak areas differed between chemical compounds, whereby the variance was close to 0 for age, only slightly higher for sex and more pronounced for group. Again, variance related to control predictors such as season of the year was considerably higher than for sex, age and group (see Table 4).

When investigating the compound-specific estimates per test predictor, the credible intervals for all compound-specific age estimates overlapped 0, although one amide, one aromatic hydrocarbone and an unidentified compound overlapped by < 20 %. Similarly, the credible intervals for all compound-specific sex estimates overlapped between the sexes, although one aldehyde, one monoterpene and 4 unidentified compounds showed only a minor overlap (i.e. central estimate for one sex outside the credible interval of the other sex, Supplement 3). Putative chemical identifications for these compounds are reported in Table S7 of Supplement 1.

The group estimates for 8 compounds did not overlap between at least 2 of the 3 groups, suggesting that the peak areas of these compounds differ distinctly between groups (Fig. 1, Supplement 3). We tentatively identified 5 of these compounds as a ketone, a carboxylic acid, an alcohol, an aromatic hydrocarbon and a catechol derivate (Table 5).

**Figure 1:**
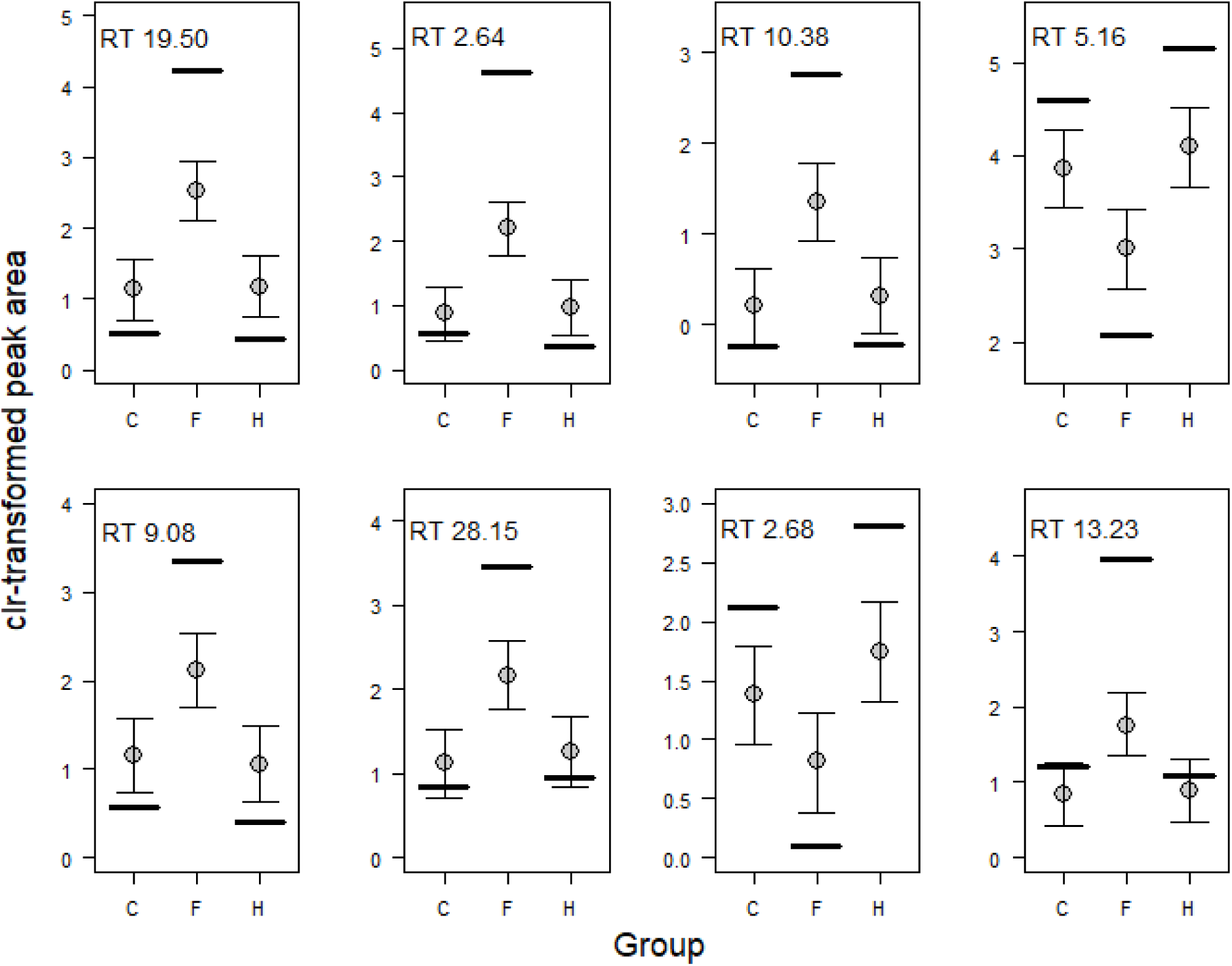
Group differences (C, F and H) in peak areas (centered log-ratio transformed) of 8 compounds with retention times in [min] as indicated in the upper left corner of each graph. Compounds are sorted by order of importance as derived from their compound-specific slope estimates. Vertical lines show mean peak areas per group, points the estimates per group (with all other predictors at their average) and whiskers the 95 % Credible Intervals.

**Table 5:**
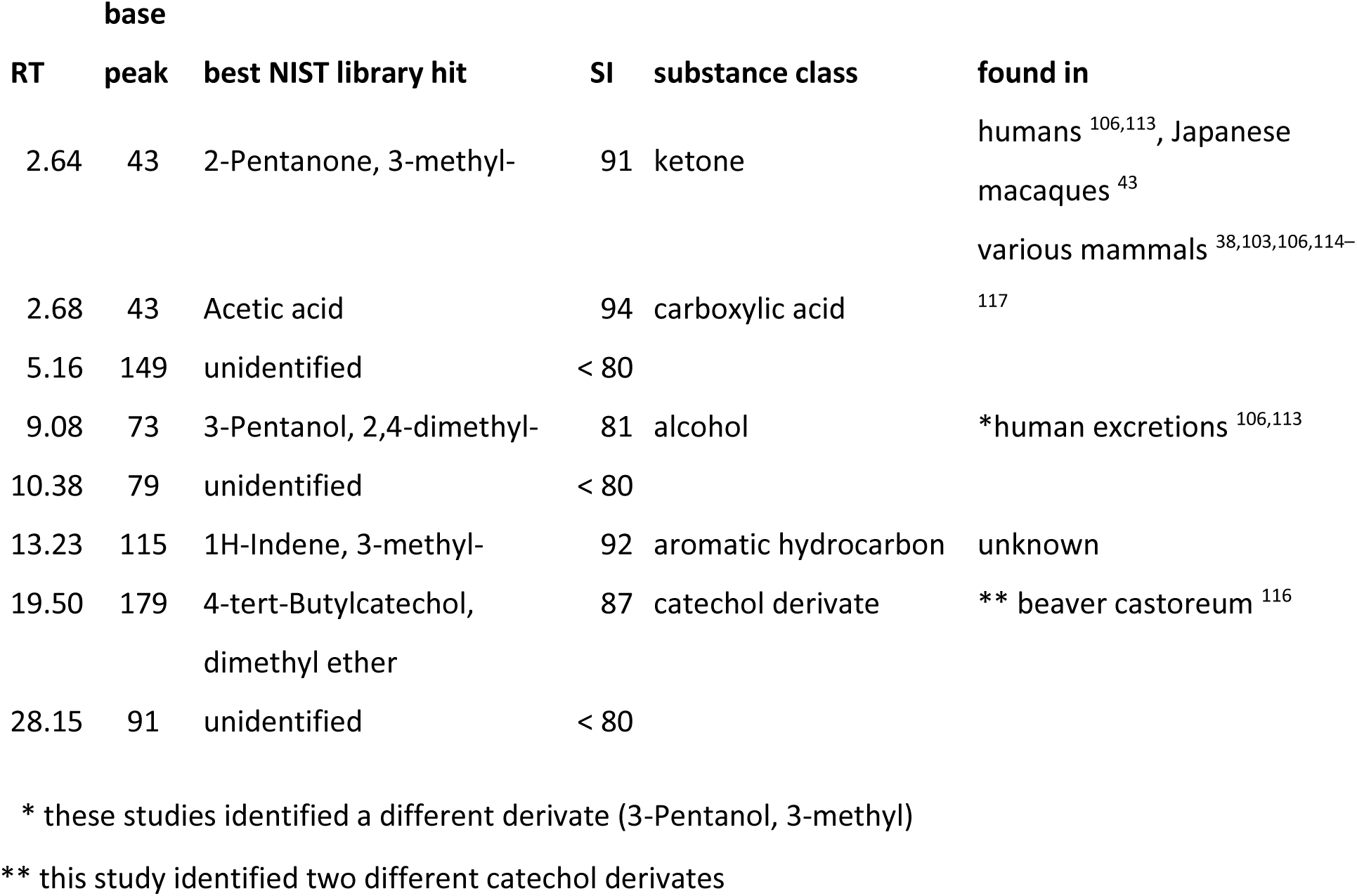
Compounds associated with group differences in chemical profiles of Barbary macaque samples. RT = retention time (in min.), base peak describes the largest mass fragment of the peak. SI = similarity index (from 0 - 100) for the match between mass spectra and their respective best hit in the NIST library.

## Discussion

In the present study, we investigated to what extent the chemical composition of Barbary macaque body odor is related to socially relevant traits and might thus have the potential to convey social information. Our analyses support the existence of individual odor signatures, while support for kin odor signatures differed between our analytical approaches. Chemical profiles were also related to group membership, but we found little to no support that rank, sex or age affect chemical similarity or detailed composition of odors emanating from the skin and fur of Barbary macaques.

Individual identity showed the strongest and most robust signature related to social information in Barbary macaque body odor. Our results are in line with the extensive evidence for individually distinct body odor across a range of mammals ^e.g. 86–88^, including primates ^e.g. 89,90,43,91,92,reviewed^ ^in^ ^85^.

While the majority of evidence thus far has come from scent-marking species, our results, along with studies in Japanese macaques ^43^ and on human axillary odor ^93^, indicate that individually distinct odors may also be present in catarrhine primates that do not show apparent scent-marking.

Individual identity explained only slightly less variance (r^2^ of 0.2) in the chemical profiles of Barbary macaques than in Japanese macaque urine (r^2^ of 0.25) or vaginal secretions (r^2^ of 0.29). Although the comparison between Japanese and Barbary macaques certainly needs to be viewed in light of different methods and a small sample size in the Japanese macaque study^43^, it suggests that odor emitted from the body surface may carry identity information of similar magnitude as potentially more intense odor sources. Furthermore, the present study shows that individually distinct information can be conveyed by the chemicals emitted from the body surface, not just by odors deposited in the environment in the form of scent marks or excretions.

The comparison of close vs. distant or non-kin based on overall chemical similarities suggested olfactory signatures of both maternal and paternal relatedness, although paternal signatures were statistically less robust. Units of close kin also showed a more similar chemical composition in owl monkeys (*Aotus nancymaae*) ^94,but^ ^see^ ^91^ and rhesus macaques ^40^. Similar to our results in Barbary macaques, kin signatures in rhesus macaques were only statistically robust among maternal, but not paternal half-siblings ^40^. These patterns suggest that similarities in body odor between maternal relatives may to some extent be mediated by frequent physical contact and/or a shared environment, which could result in an exchange of odor-producing microbes, as e.g. suggested for group odor signatures in hyenas ^95^. However, when treating relatedness as a continuous variable as well as accounting for more other potential sources of variation in body odor, we found no support whatsoever for relatedness predicting chemical similarity between individual odors. Possibly, chemical profiles of surface body odor allow only a coarse, but no fine-scaled differentiation of relatedness. Alternatively, what appeared to be kin signatures in the perMANOVAs may actually have been variation related to other predictors that could not be as accurately controlled for in these analyses. Furthermore, kin odor signatures may only be robustly expressed in certain contexts or seasons, when kin discrimination is particularly important. For instance, kin signatures in glandular secretions of ring-tailed lemurs (*Lemur catta*) were robustly expressed only in the breeding season, when they may aid in inbreeding avoidance ^96,97^, but we did not collect odor samples during the mating season for logistic reasons.

The lack of robust chemical information about kinship involving the paternal line (at least outside the mating season) is especially interesting with respect to the triadic interactions and care that Barbary macaque males frequently engage in with infants. Particularly, male-infant interactions have been suggested to be independent of paternity to the respective infant ^53^, which may be due to low phenotypic similarity between father and offspring when the offspring is still in its infancy. Studies in rhesus macaques testing exclusively adult subjects suggest that unfamiliar, paternal kin discriminate relatives based on visual or acoustic phenotypic cues ^9,10^. Moreover, humans rating the facial similarity of rhesus macaque parents and offspring performed above chance level ^98^ and better if the offspring were older ^99^. The paternal kin units sampled in the present study indeed comprised cases in which the offspring was still rather young (2 years old), but in half of the cases all members of a paternal kin unit were reproductively mature (i.e. 4 years or older). Hence, the chemical data presented here do not point towards olfactory signatures alone being robust enough to allow Barbary macaque males to reliably distinguish between own and unrelated offspring.

Regardless of the analytical approach, we detected no robust support for chemical profiles being related to dominance classes or rank. Although we found a significant effect of dominance class on the similarity between chemical profiles, the association was very weak, and ranks across the entire dominance hierarchy were associated with neither chemical similarity nor the abundance of specific compounds. Dominance-related odors have been described in various mammals ^e.g. 61,reviewed^ ^in^ ^92^, including male mandrills ^37,38^, but were not detected in ring-tailed lemurs ^92^ or female mandrills ^37^. In female rhesus macaques, individual rank had no effect on overall chemical similarities, but affected the relative abundance of certain compounds ^40^. As pointed out by Smith ^85^, olfactory communication of dominance requires either a specific dominance-related odor or olfactory individual signatures that can be associated with other dominance-related information. Evidence for the latter comes from a study in ring-tailed lemurs, in which males discriminated between the odors of dominant and subordinate individuals, but only if they were familiar with the odor donors ^21^. Hence, while our results provide no robust evidence for dominance-related body odor in either male or female Barbary macaques, the detected individual odor signatures certainly open the possibility for a more indirect olfactory communication of dominance among familiar individuals. Furthermore, our results imply that Barbary macaques of similar rank either shared spaces less than we had assumed, or that sharing similar spaces had no detectably effect on chemical profiles. The latter seems contradictory to our suggestion that a shared environment may have contributed to the more robust odor signatures among close maternal than paternal kin. However, a higher degree of spatial proximity and physical contact among close maternal kin than among unrelated individuals of similar rank could lead to kin, but not rank signatures to be affected by the environment.

Based on behavioral and/or chemical evidence, the odors of numerous primates and other mammals convey sex-specific ^18,40,91,100,101^ and age-specific information ^24,86,94,100,102,103^, although this need not always be the case ^102,also^ ^see^ ^30^. Age, for instance, was related to chemical profiles in one subset of individuals in rhesus macaques, but not in another ^40^. In the present study, sex and age, although significantly associated with chemical similarities, explained very little variation in chemical profiles, and none of the chemical compounds distinctly differed in abundance between sexes or showed consistent changes with age. It is possible that sex and age information may actually be more pronounced in a different context (e.g. mating), in other odor sources (e.g. urine) or other body parts (e.g. the ano-genital region), all of which, along with differences in sampling and analytical methods, may affect the expression of chemical signatures or our ability to detect them ^30,43,see^ ^97,104^.

Furthermore, sex differences may have been attenuated in the study population due to the use of contraceptives in 60 % of the adult females (see below).

Group membership explained about 5 % of the variance in chemical similarities and affected the relative abundance of 8 compounds. For most of these compounds, abundances were consistently higher or lower in group F, but similar in the other two groups. Group odor signatures have been described in a range of group-living mammals, including rhesus macaques ^39,40^, mandrills ^38^ and chimpanzees ^41^, with explanations for the origins of these differences including the environment, diet and microbial differences between groups ^91,95,101^. In species living in family groups, group odor signatures may further reflect relatedness among individuals (e.g. in owl monkeys ^94^). Relatedness is an unlikely explanation for the group differences observed in the present study, because all groups comprised multiple matrilines and average relatedness among genotyped focal animals was low both within (r = 0.04) and between groups (r = 0.02, data in Supplement dyadic relatedness data). The three groups also received the same supplemental food and had spatially (but not temporally) overlapping home ranges, but there are subtle differences in the vegetation in different areas of the park that could potentially affect natural food sources as well as background volatiles or substances the animals get into contact with. Several of the tentatively identified compounds could fit such an explanation. 3-methyl-2-pentanone and 2,4-dimethyl-3-pentanol, for instance, could be microbial or plant products metabolized by the mammalian body ^see^ ^105^ and similar compounds have been found in human feces before ^106^. Furthermore, indenes have been described in plants ^e.g. 107^, although we could not find a source specifically for 3-methyl-1H-indene, which was the best mass spectral library match for the compound at RT 13.23 min. Alternatively, group odor differences could be related to demographical and social differences between groups. The focal animals of group F comprised a slightly higher proportion of females than groups C and H (57 % vs. 43 % and 32 %) and were marginally younger (average age in group F 9.2 years vs. 11.7 and 11.3 yrs in groups C and H, see Supplement 1 Tab. S2). However, because sex and age effects on body odor were very small (see above) and statistically accounted for, they are unlikely to contribute much to the observed group differences. Group F also is the largest and highest-ranking of the three groups, which could affect stress levels or other physiological parameters ^see^ ^108^ and thereby, body odor (reviewed in ^109^), but we have no physiological data available to further assess this possibility.

Main assets of our study certainly are the large numbers of samples and focal individuals, the semi-natural environment and natural social conditions. We also were able to collect all samples non-invasively, thereby reducing the chance that odors were affected by sampling-related stress. At the same time, the study faced several limitations, some of which we were able to account for during data analysis, and others not. For instance, we collected samples from different body parts that we could approach most easily, i.e. primarily the shoulders, flanks and thighs, and controlled for the variation in sampled body regions during statistical analysis. However, it is difficult to assess how relevant these body parts actually are for day-to-day information exchange. Recent observational data in the study population suggest that sniffs directed at conspecifics frequently target the ano-genital region of adult females in a mating context ^60^. The ano-genital region might thus be a more appropriate sampling region at least for certain types of individuals and contexts, but would have been feasible to sample only in a subset of very well-habituated females at frequent intervals. As the social information detectable in animal odors may differ between odor sources, even within species ^see^ ^30,43^, it is possible that results would have differed if we had sampled other body regions or odor sources such as urine.

Our data collection periods spanned much of the year, but for logistic reasons we did not collect data during most of the park’s winter break, which corresponds to the peak of the mating season in the study population. Indeed, the season of the year did affect chemical profiles, even more so than the socially relevant variables investigated. In particular, seasonal variation may have affected the volatile background in the environment, which we could at least in part account for by using environmental control samples. Furthermore, aspects such as reproductive physiology, diet, ambient temperature or state of the fur (winter vs. summer coat) may additionally affect the chemical compounds emitted by the body and how readily they become available to recipients (or sampling tube). Such aspects are likely to be at the base of the detected seasonal changes in chemical profiles, and we statistically accounted for them in our analyses of socially related olfactory signatures by using season of the year as a control variable. What remains open is whether any of the investigated effects might have been more pronounced during the mating season, as shown, e.g., for kin signatures in ring-tailed lemurs ^96,97^.

Another aspect to consider is that the size of the study population is managed via hormonal contraception, and 60 % of the sampled females received contraception during part or all of the study period. Studies on the effects of hormonal contraception on primate body odor are scarce and contradicting: hormonal contraception using medroxyprogesterone acetate has previously been shown to alter the chemical composition of female odors in ring-tailed lemurs ^110^ but not in owl monkeys ^91^. Male mandrills showed more flehmen responses towards odor samples from cycling females than from females contracepted with etonogestrel ^111^, which is the active ingredient of the implants used in the studied Barbary macaques. However, while contraception also tended to modulate age-specific differences in ano-genital sniffs by male Barbary macaques, it did not affect subsequent sexual behavior in our studied population ^60^. As we did not assess individual attributes immediately related to mating and female fertility, contraception might be considered as a source of noise in the data. Unlike several other sources of noise, however, we could not account for contraception statistically given that it applied only to a minority of samples and individuals. Overall, it is thus possible that certain signatures, in particular those of sex, might have been more pronounced in a sample without females receiving contraception.

Our results also suggest some variation in chemical profiles related to sampling and GC-MS measurement, whereby different variables affected different aspects of the composition: chemical similarities between samples were greater if samples were collected closer in time, and to a lesser extent also when analyzed within the same GC-MS batch. In contrast, variables such as the sampling distance, sampling volume or the sampled body part had little effect on chemical similarities of samples, but appeared to have some effect on the relative abundance of certain compounds.

However, while we were unable to eliminate these sources of variation in the study design, we statistically accounted for them in all mixed model analyses.

Finally, results need to be put into perspective of what we can measure vs. what animals can perceive. Statistically detectable associations between chemical profiles and socially relevant traits show the potential for olfactory information transfer, but we do not know if or how animals use this information. Animal sensory systems have evolved for the detection of relevant information and may thus be superior to even the most modern of our analytical techniques, at least for certain compounds (reviewed in ^112^). Accordingly, it may be possible that animals are able to pick up upon even very subtle differences in odors, or differences that our analyses are unable to pick up at all, especially as we are unlikely to sample and detect all chemical substances emitted by an animal’s body across the possible ranges of volatility, polarity etc. ^104^ despite the constant improvements in sampling and analytical methods. Hence, future studies should combine chemical data with (experimental) assessments of behavioral and physiological reactions towards conspecific odors across a range of contexts to obtain a detailed understanding of the complex role of olfaction in primate social interactions. Particularly under natural and semi-natural conditions, controlled presentations of conspecific odors are challenging to implement ^e.g. 55^, as stimuli need to be presented in a manner that is salient and realistic enough to elicit interest and closer inspection without leading to quick habituation. Promising next steps would be multi-modal experiments and experimental manipulation of *in vivo* odors to investigate how olfactory discrimination of social information falls into place between the other senses, i.e. whether macaques are able to discriminate such information based on odors alone, and/or if body odor signatures improve the reliability of other sensory cues.

In conclusion, our results add to the limited evidence that socially relevant information may not only be encoded in odor secretions and excretions, but also in the volatile and semi-volatile components emanating from the body surface of primates. Based on chemical profiles alone, i.e. without knowing what the monkeys can make of it, data suggest that Barbary macaque body odor could directly contribute to recognizing individuals and groups, while body odor signatures may not be pronounced or robust enough to allow differentiating kinship, rank, sex or age of conspecifics from surface body odor alone. However, given the central role of individual recognition in forming and maintaining social relationships, surface body odor could potentially still contribute to managing social relationships, if monkeys associate social knowledge obtained via other sensory modalities or direct social interactions with individual odor signatures.

## Acknowledgements

We thank Roland & Mamisolo Hilgartner and the staff of the Affenberg Salem for continuous support and Ellen Merz for permission to conduct this study. We further thank Roger Mundry for an introduction to multi-membership models and help with the centred log-ratio transformation.

Hendrikje Westphal established the methods for genetic paternity analyses and helped with faecal sample collection. Stefanie Bley did the allele calling and managed the paternity analyses. Antonia Krüger and Anja Schiffner conducted DNA extractions and PCR analyses, and helped with GC-MS measurements and TD tube cleaning. Annika Freudiger helped with the implementation of the TRACE software. Ramona Oehme and Susan Billig in the mass spectrometry core facility at the Faculty of Chemistry and Mineralogy at Leipzig University provided technical support in GC-MS maintenance.

This work was supported by the German Research Foundation, DFG (grant no. SCHL 2011/2-1 awarded to B. M. Weiß). The GC-MS instrument was funded by the European Fund for Regional Structure Development, EFRE (“Europe funds Saxony”, grant #100195810 to A. Widdig).

