## Supplement 1 for "Chemical signatures of social information in Barbary macaques"

Supplemental methods and results

*Details on paternity analysis*

The 144 sampled individuals included 28 offspring from cohorts 2017-2019 (for 22 of which we also collected odor samples), their mothers and potential sires. Because Barbary macaques may also reproduce with extra-group partners ^1^, we considered all males in the population that were alive and at least 4 years of age at the onset of the respective mating season as potential sires. For animals born in cohort 2017 we were able to obtain samples for 96 %, and for cohorts 2018-2019, 97 % of all potential sires.

For 27 individuals blood samples had been collected during medical interventions prior to the onset of the study. Similarly, 7 tissue samples had previously been collected during medical intervention or from freshly deceased bodies. In recent years, hair samples have been routinely collected during tattooing of yearlings, so that hair samples were available for 68 of the individuals. To obtain DNA from 41 potential sires and one mother for whom no other sample types were available, we collected three fecal samples per individual immediately after defecation. Fecal samples were stored with the two-step alcohol-silica storage protocol ^2^. DNA was extracted using DNEasy Blood & Tissue kit (Qiagen) for blood and tissue, and the First-DNA all-tissue DNA-Kit (GEN-IAL) for hair and feces.

We genotyped the DNA samples using a panel of up to 23 polymorphic microsatellite markers previously established for the study population ^3^, whereby we eventually focused on the 16 markers with the lowest dropout rate, that were most reliable, scorable and combinable in polymerase chain reactions (PCR, see Table S1). To increase the amount of host DNA in the samples we used the two-step multiplex approach for PCR amplification ^4^ by running a first PCR with all markers simultaneously, followed by a second set of PCRs containing diluted multiplex products as amplification templates and 4 – 6 fluorescently labelled primers with different allele ranges. PCRs were conducted in a Mastercycler® vapo.protect thermal cycler (Eppendorf, Hamburg, Germany) following the protocols of Westphal ^3^. We conducted 2 independent sets of PCRs for each of the blood and tissue samples, and 3 for each of the hair and fecal samples. Due to the small amount of DNA and a high level of allelic dropouts in fecal samples ^5^, we analyzed two independent fecal samples per individual ^e.g. 6^.

We performed fragment analysis on an *ABI 3730* sequencer. For allele calling we used the Peak Scanner 2 software (Applied Biosystems®). For all sample types we confirmed heterozygous genotypes with two amplifications. For blood and tissue samples we confirmed homozygous genotypes with two, for hair samples with three and for fecal samples with four amplifications. In case of inconsistent results an additional sample of the respective individual was analyzed.

For 27 of the 28 offspring born 2017 – 2019 the observed mothers showed no mismatches to their offspring. For one offspring we could not confirm the observed mother genetically, probably due to poor sample quality. In this case we considered the observed mother to be the biological mother, but excluded the offspring from paternity analysis. We could also genetically confirm the observed mother for 38 females and males initially sampled as mothers or potential sires.

We used confirmed maternities in the paternity analysis, applying a combination of exclusion and likelihood methods for mother-offspring-putative father trios with the programs Findsire and Cervus following established criteria ^e.g. 6,7^. In particular, we used the following exclusion criteria for paternity assignment with Findsire: the strict criterion was met if a sire showed no mismatches and the next potential sire at least two and the relaxed criterion if a sire showed no mismatches and the next potential sire one mismatch. Any cases in which all potential sires had at least one mismatch were considered unsolved. Cases in which two potential sires showed no mismatch were considered a tie. Ties were also considered unsolved unless one of the potential sires was assigned a likelihood of 95% in Cervus, in which case the sire designated by Cervus was accepted as the sire. In case of inconsistent assignments by Findsire and Cervus, paternity was considered unsolved.

We were able to assign fathers to 23 of the 28 offspring from cohorts 2017-2019 with high certainty. Furthermore, we accepted fathers assigned with high confidence for 18 individuals initially sampled as mothers or potential sires in the present study and born between 2011 and 2016. For these cohorts, 58-87 % of potential sires were genotyped. We did not accept fathers assigned to older individuals due to the increasingly lower proportion of potential sires sampled.

As input we used genetically confirmed mothers whenever available, and observed mothers (based on observational records from the field) for all remaining subjects, as well as all available paternity data. Given that mother-offspring dyads are available from long-term observational records, our value of pedigree relatedness for maternal relatedness can be considered as highly precise, reflecting close maternal (r > 0.125) but also distant maternal kin (r ≤ 0.125), while our values of pedigree relatedness for the paternal line are likely underestimated in this analysis.

Table S1: Final microsatellite marker set used for parentage analysis. Primers in the same set were combined in the second PCR reaction.

| **Primer** | **tandem repeat** | **labeled** | **allele size range** | **primer set** |
| --- | --- | --- | --- | --- |
| D10s1432 | tetra | hex | 140-164 | 1 |
| D1s548 | tetra | hex | 184-208 | 1 |
| D2s1333 | tetra | hex | 290-306 | 1 |
| D12s67 | tetra | fam | 126-142 | 1 |
| D6s493 | tetra | fam | 154-166 | 1 |
| D13s765 | tetra | fam | 177-194 | 1 |
| D10s611 | tetra | hex | 157-185 | 2 |
| D19s245 | tetra | hex | 249-333 | 2 |
| D19s582 | tetra | fam | 131-143 | 2 |
| D4s243 | tetra | fam | 191-203 | 2 |
| D7s2204 | tetra | ned | 227-243 | 2 |
| D5s1467 | tetra | hex | 192-200 | 3 |
| D5s1466 | tetra | hex | 273-329 | 3 |
| D7s503 | di | fam | 142-164 | 3 |
| D1s518 | tetra | fam | 186-206 | 3 |
| D11s925 | di | ned | 189-205 | 3 |

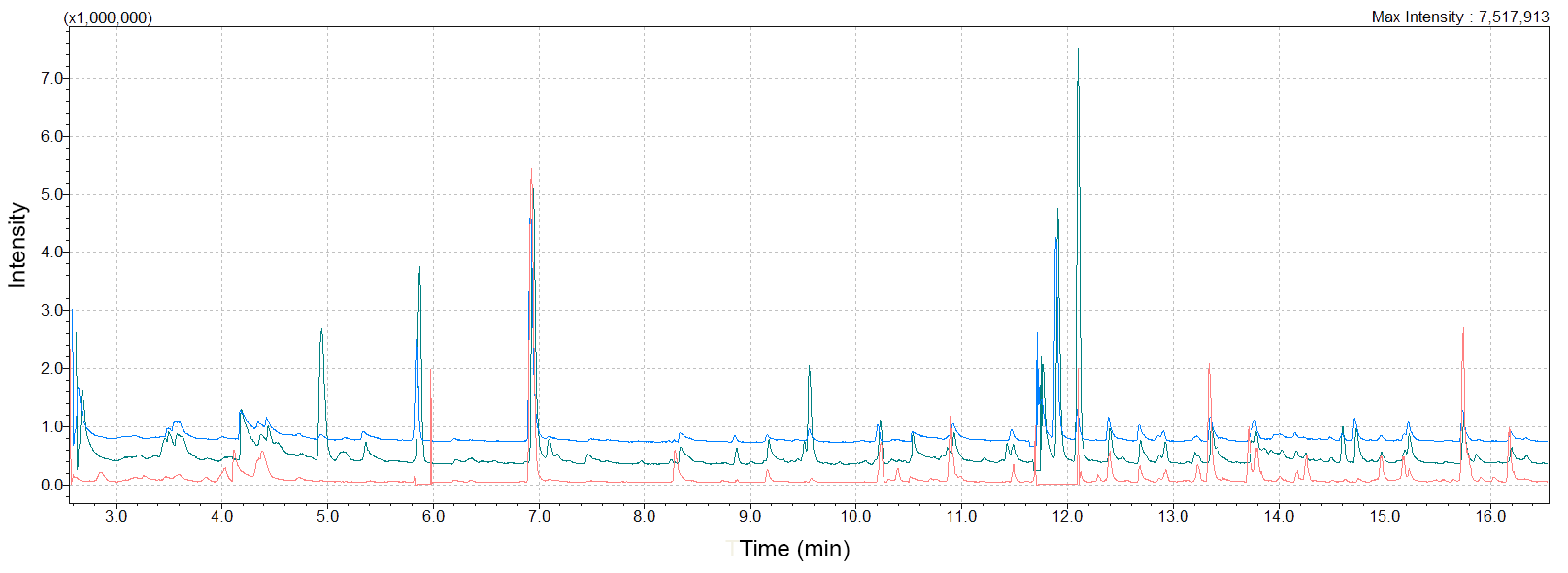

*Figure S1*: Chromatograms of Barbary macaque body odor samples from two different females (blue and green lines) and a blank control sample (orange line) over the volatile range. Base lines were slightly shifted along the y-axis for better visibility of the individual lines.

Table S2: List of focal individuals, their identity (IDcode), sex, year of birth (cohort) and group membership. "contraception" denotes for all adult females if the individual received hormonal contraception during the study period. "rank_cat", "matkin_unit" and "patkin_unit" denote the dominance category and family unit membership for those of the focal individuals used in the respective analysis.

| IDcode | sex | cohort | group | contraception | rank_cat | matkin_unit | patkin_unit |
| --- | --- | --- | --- | --- | --- | --- | --- |
| ID01 | male | 2017 | H | NA | NA | m7 | p5 |
| ID02 | male | 2009 | C | NA | NA | NA | NA |
| ID03 | male | 2010 | H | NA | low | NA | NA |
| ID04 | female | 2018 | F | no | NA | m10 | p9 |
| ID05 | male | 2004 | C | NA | high | NA | p4 |
| ID06 | male | 2007 | H | NA | high | NA | NA |
| ID07 | female | 2017 | F | no/yes | low | NA | NA |
| ID08 | male | 2006 | H | NA | NA | NA | p7 |
| ID09 | male | 2009 | H | NA | low | m6 | NA |
| ID10 | female | 2012 | H | yes | NA | NA | NA |
| ID11 | male | 2009 | C | NA | NA | NA | p3 |
| ID12 | female | 2011 | C | yes | high | m11 | NA |
| ID13 | female | 2012 | H | no | high | m7 | NA |
| ID14 | male | 2011 | F | NA | NA | NA | p2 |
| ID15 | female | 2017 | F | no | NA | NA | p1 |
| ID16 | female | 2002 | H | no | NA | NA | NA |
| ID17 | female | 2011 | C | yes | NA | m4 | NA |
| ID18 | female | 2017 | C | no | NA | m4 | p12 |
| ID19 | female | 2009 | F | yes | high | NA | NA |
| ID20 | female | 2012 | F | yes | low | m2 | NA |
| ID21 | male | 2008 | H | NA | NA | NA | NA |
| ID22 | male | 2004 | C | NA | NA | NA | p12 |
| ID23 | male | 2017 | F | NA | NA | m5 | p2 |
| ID24 | female | 2012 | F | yes | NA | m5 | NA |
| ID25 | male | 1999 | H | NA | high | NA | NA |
| ID26 | female | 2018 | F | no | NA | m8 | NA |
| ID27 | male | 2000 | F | NA | NA | NA | p1 |
| ID28 | male | 2007 | C | NA | low | NA | NA |
| ID29 | female | 2019 | C | no | NA | m12 | p12 |
| ID30 | male | 2009 | C | NA | high | NA | NA |
| ID31 | male | 2006 | F | NA | NA | NA | p11 |
| ID32 | male | 2004 | H | NA | NA | NA | p5 |
| ID33 | male | 2019 | F | NA | NA | m9 | p6 |
| ID34 | female | 2006 | F | yes | high | m1 | NA |
| ID35 | male | 2008 | F | NA | high | NA | p10 |
| ID36 | male | 2013 | F | NA | low | NA | p6 |
| ID37 | male | 2009 | C | NA | low | NA | p8 |
| ID38 | female | 2018 | H | no | NA | NA | p5 |
| ID39 | female | 2019 | C | no | NA | m3 | p3 |
| ID40 | male | 2013 | C | NA | low | NA | NA |
| ID41 | male | 2018 | H | NA | NA | m6 | NA |
| ID42 | female | 2007 | F | yes | NA | NA | NA |
| ID43 | male | 2003 | C | NA | high | NA | NA |
| ID44 | male | 2008 | F | NA | high | NA | NA |
| ID45 | female | 2014 | C | yes | high | m12 | p12 |
| ID46 | female | 2005 | H | yes | low | m6 | NA |
| ID47 | male | 2017 | H | NA | NA | NA | NA |
| ID48 | female | 2008 | C | yes/no | low | m10 | NA |
| ID49 | male | 2019 | F | NA | NA | m1 | p11 |
| ID50 | male | 2008 | C | NA | low | NA | NA |
| ID51 | female | 2014 | C | yes | high | m3 | NA |
| ID52 | male | 1995 | C | NA | NA | NA | NA |
| ID53 | male | 1999 | F | NA | NA | NA | NA |
| ID54 | male | 2007 | H | NA | NA | NA | NA |
| ID55 | female | 2013 | F | yes | low | m10 | NA |
| ID56 | male | 2018 | C | NA | NA | m11 | p4 |
| ID57 | female | 2011 | F | yes | high | m1 | NA |
| ID58 | female | 2015 | F | yes | high | m1 | p8 |
| ID59 | female | 2017 | F | no | NA | NA | p10 |
| ID60 | female | 2005 | F | yes | NA | NA | NA |
| ID61 | male | 2017 | C | NA | NA | m11 | p3 |
| ID62 | female | 2014 | C | no | low | NA | NA |
| ID63 | male | 2018 | H | NA | NA | NA | p7 |
| ID64 | female | 2017 | F | no | low | m2 | p2 |
| ID65 | male | 2009 | F | NA | low | NA | NA |
| ID66 | male | 2018 | H | NA | NA | m7 | NA |
| ID67 | male | 2012 | F | NA | high | NA | NA |
| ID68 | female | 2013 | F | yes | NA | m8 | NA |
| ID69 | female | 2008 | H | yes | high | m7 | NA |
| ID70 | female | 2014 | F | yes | low | m9 | NA |
| ID71 | male | 2013 | F | NA | low | NA | p9 |
| ID72 | female | 2017 | C | no | NA | NA | NA |

*Detailed results tables*

Table S3: Detailed results of perMANOVAs conducted with data subsets for ID, kinship and dominance class.

| **subset** | **term** | **Df** | **SumOfSqs** | **R2** | **F** | **P** |
| --- | --- | --- | --- | --- | --- | --- |
| ID | ID | 21 | 12.084 | 0.203 | 3.838 | 0.001 |
|  | sex | 0 | 0.000 | 0 | Inf |  |
|  | age | 1 | 7.620 | 0.128 | 50.823 | 0.001 |
|  | group | 1 | 0.072 | 0.001 | 0.478 | 0.989 |
|  | season | 2 | 8.085 | 0.136 | 26.961 | 0.001 |
|  | residual | 214 | 32.087 | 0.538 |  |  |
|  | total | 241 | 59.628 | 1 |  |  |
| maternal kin | kin unit | 11 | 5.27 | 0.099 | 2.537 | 0.001 |
|  | sex | 1 | 0.276 | 0.005 | 1.461 | 0.001 |
|  | season | 2 | 7.74 | 0.145 | 20.493 | 0.001 |
|  | age | 1 | 0.47 | 0.009 | 2.488 | 0.001 |
|  | residual | 204 | 38.523 | 0.72 |  |  |
|  | total | 219 | 53.479 | 1 |  |  |
| paternal kin | kin unit | 11 | 5.877 | 0.132 | 2.772 | 0.067 |
|  | sex | 1 | 0.201 | 0.005 | 1.043 | 0.001 |
|  | season | 2 | 4.956 | 0.111 | 12.855 | 0.001 |
|  | age | 1 | 0.340 | 0.008 | 1.761 | 0.790 |
|  | residual | 163 | 31.419 | 0.706 |  |  |
|  | total | 178 | 44.528 | 1 |  |  |
| dominance class | sex | 1 | 1.254 | 0.019 | 6.535 | 0.001 |
|  | dominance class | 1 | 0.169 | 0.003 | 0.879 | 0.008 |
|  | age | 1 | 0.769 | 0.012 | 4.007 | 0.001 |
|  | group | 2 | 3.063 | 0.047 | 7.982 | 0.026 |
|  | season | 2 | 8.856 | 0.136 | 23.080 | 0.001 |
|  | residual | 252 | 48.345 | 0.742 |  |  |
|  | total | 259 | 65.153 | 1 |  |  |

Table S4: Estimates for all fixed and random terms from the multi-membership model on genetic relatedness rank. CrI = Credible Interval. ESS = effective sample size. “d_xxx” denotes delta, i.e. the difference of the respective term between the two samples in a given dyad.

| **fixed term** | **Estimate** | **Est.Error** | **2.5% CrI** | **97.5% CrI** | **Rhat** | **bulk ESS** | **tail ESS** |
| --- | --- | --- | --- | --- | --- | --- | --- |
| Intercept | 0.142 | 0.026 | 0.089 | 0.194 | 1.002 | 1592 | 2884 |
| maternal r | 0.000 | 0.001 | -0.002 | 0.002 | 1.000 | 9718 | 7456 |
| paternal/mixed r | -0.002 | 0.002 | -0.005 | 0.001 | 1.000 | 6365 | 5522 |
| d_elo score | 0.000 | 0.001 | -0.002 | 0.002 | 1.000 | 10551 | 7505 |
| d_sex (same) | -0.002 | 0.002 | -0.005 | 0.002 | 1.000 | 9026 | 7614 |
| d_age | -0.001 | 0.002 | -0.004 | 0.002 | 1.000 | 6827 | 6459 |
| d_group (same) | 0.012 | 0.006 | 0.001 | 0.023 | 1.001 | 2163 | 3956 |
| d_season (same) | -0.024 | 0.012 | -0.048 | -0.001 | 1.003 | 562 | 1399 |
| d_date | -0.320 | 0.009 | -0.339 | -0.301 | 1.007 | 528 | 944 |
| d_time | 0.001 | 0.001 | -0.002 | 0.003 | 1.000 | 4987 | 6121 |
| d_bodyregion (same) | 0.003 | 0.003 | -0.003 | 0.008 | 1.001 | 4418 | 5675 |
| d_s.distance | 0.004 | 0.002 | 0.000 | 0.009 | 1.001 | 3783 | 5261 |
| d_s.volume | 0.007 | 0.005 | -0.004 | 0.016 | 1.003 | 984 | 1902 |
| d_subsamples | -0.007 | 0.002 | -0.011 | -0.003 | 1.001 | 4902 | 6101 |
| d_batch (same) | 0.199 | 0.015 | 0.169 | 0.230 | 1.001 | 1051 | 1941 |
| **random term** | **Estimate** | **Est.Error** | **2.5% CrI** | **97.5% CrI** | **Rhat** | **bulk ESS** | **tail ESS** |
| sample (Intercept) | 0.399 | 0.014 | 0.372 | 0.429 | 1.003 | 1074 | 2166 |
| ID (Intercept) | 0.149 | 0.028 | 0.096 | 0.204 | 1.003 | 471 | 973 |
| ID (maternal r) | 0.003 | 0.002 | 0.000 | 0.007 | 1.001 | 1781 | 3738 |
| ID (paternal/mixed r) | 0.007 | 0.002 | 0.003 | 0.012 | 1.002 | 1739 | 1761 |
| ID (d_elo score) | 0.003 | 0.002 | 0.000 | 0.007 | 1.000 | 1876 | 3449 |
| ID (d_sex) | 0.003 | 0.003 | 0.000 | 0.009 | 1.000 | 2608 | 3813 |
| ID (d_age) | 0.009 | 0.002 | 0.005 | 0.013 | 1.000 | 2379 | 2364 |
| ID (d_group) | 0.043 | 0.005 | 0.035 | 0.053 | 1.000 | 3230 | 5042 |
| ID (d_season) | 0.094 | 0.009 | 0.078 | 0.113 | 1.002 | 1607 | 3186 |
| ID (d_date) | 0.081 | 0.007 | 0.068 | 0.097 | 1.002 | 913 | 2059 |
| ID (d_time) | 0.009 | 0.001 | 0.006 | 0.012 | 1.001 | 3418 | 5151 |
| ID (d_bodyregion) | 0.019 | 0.003 | 0.014 | 0.026 | 1.000 | 3248 | 4049 |
| ID (d_s.distance) | 0.016 | 0.002 | 0.012 | 0.020 | 1.000 | 3528 | 5360 |
| ID (d_s.volume) | 0.041 | 0.004 | 0.034 | 0.049 | 1.000 | 1500 | 3193 |
| ID (d_subsamples) | 0.013 | 0.002 | 0.010 | 0.018 | 1.000 | 3736 | 5669 |
| ID (d_batch) | 0.124 | 0.012 | 0.103 | 0.149 | 1.001 | 2176 | 4369 |
| dyad (Intercept) | 0.019 | 0.001 | 0.016 | 0.021 | 1.001 | 1690 | 3502 |
| dyad (d_season) | 0.040 | 0.002 | 0.036 | 0.044 | 1.000 | 2014 | 4321 |
| dyad (d_date) | 0.005 | 0.002 | 0.000 | 0.009 | 1.012 | 462 | 1413 |
| dyad (d_time) | 0.007 | 0.002 | 0.003 | 0.009 | 1.003 | 902 | 634 |
| dyad (d_bodyregion) | 0.029 | 0.002 | 0.025 | 0.032 | 1.001 | 2629 | 4323 |
| dyad (d_s.distance) | 0.017 | 0.001 | 0.015 | 0.019 | 1.001 | 2302 | 4114 |
| dyad (d_s.volume) | 0.012 | 0.001 | 0.009 | 0.014 | 1.002 | 1642 | 2844 |
| dyad (d_subsamples) | 0.016 | 0.001 | 0.014 | 0.018 | 1.002 | 2199 | 4233 |
| dyad (d_batch) | 0.087 | 0.003 | 0.081 | 0.093 | 1.000 | 2602 | 5162 |

Table S5: Estimates for all fixed and random terms from the multi-membership model on rank. CrI = Credible Interval. ESS = effective sample size. “d_xxx” denotes delta, i.e. the difference of the respective term between the two samples in a given dyad.

| **fixed term** | **Estimate** | **Est.Error** | **2.5% CrI** | **97.5% CrI** | **Rhat** | **bulk ESS** | **tail ESS** |
| --- | --- | --- | --- | --- | --- | --- | --- |
| Intercept | 0.154 | 0.039 | 0.076 | 0.232 | 1.001 | 2680 | 5074 |
| d_elo score | 0.000 | 0.001 | -0.003 | 0.003 | 1.000 | 19102 | 9841 |
| total r | -0.002 | 0.001 | -0.005 | 0.001 | 1.000 | 18584 | 8977 |
| sex (male) | -0.023 | 0.053 | -0.126 | 0.080 | 1.001 | 2879 | 5192 |
| d_age | 0.001 | 0.002 | -0.003 | 0.005 | 1.000 | 14446 | 9510 |
| d_group (same) | 0.016 | 0.007 | 0.002 | 0.030 | 1.000 | 5497 | 7723 |
| d_season (same) | -0.027 | 0.011 | -0.048 | -0.006 | 1.002 | 2736 | 4694 |
| d_date | -0.325 | 0.010 | -0.346 | -0.305 | 1.002 | 1088 | 2508 |
| d_time | 0.001 | 0.001 | -0.002 | 0.003 | 1.000 | 15821 | 9039 |
| d_bodyregion (same) | 0.004 | 0.003 | -0.002 | 0.011 | 1.000 | 11941 | 8932 |
| d_s.distance | 0.005 | 0.002 | 0.001 | 0.010 | 1.001 | 11074 | 9460 |
| d_s.volume | 0.008 | 0.005 | -0.003 | 0.019 | 1.001 | 2684 | 3564 |
| d_subsamples | -0.007 | 0.002 | -0.011 | -0.003 | 1.000 | 15094 | 10231 |
| d_batch (same) | 0.184 | 0.018 | 0.148 | 0.219 | 1.001 | 2652 | 5513 |
| **random term** | **Estimate** | **Est.Error** | **2.5% CrI** | **97.5% CrI** | **Rhat** | **bulk ESS** | **tail ESS** |
| sample (Intercept) | 0.395 | 0.014 | 0.368 | 0.424 | 1.002 | 2096 | 4665 |
| ID (Intercept) | 0.151 | 0.028 | 0.099 | 0.208 | 1.006 | 891 | 1598 |
| ID (d_elo score) | 0.003 | 0.002 | 0.000 | 0.008 | 1.002 | 3549 | 6077 |
| ID (total r) | 0.003 | 0.002 | 0.000 | 0.009 | 1.001 | 3539 | 5713 |
| ID (d_age) | 0.010 | 0.003 | 0.004 | 0.016 | 1.001 | 2014 | 1208 |
| ID (d_group) | 0.053 | 0.007 | 0.041 | 0.067 | 1.001 | 5572 | 8358 |
| ID (d_season) | 0.089 | 0.009 | 0.074 | 0.108 | 1.001 | 4696 | 7678 |
| ID (d_date) | 0.086 | 0.008 | 0.072 | 0.103 | 1.001 | 2359 | 4170 |
| ID (d_time) | 0.007 | 0.002 | 0.002 | 0.011 | 1.001 | 2548 | 2941 |
| ID (d_bodyregion) | 0.021 | 0.004 | 0.013 | 0.030 | 1.000 | 4347 | 5327 |
| ID (d_s.distance) | 0.016 | 0.003 | 0.011 | 0.022 | 1.001 | 5362 | 7222 |
| ID (d_s.volume) | 0.042 | 0.004 | 0.035 | 0.051 | 1.001 | 5052 | 7551 |
| ID (d_subsamples) | 0.011 | 0.003 | 0.005 | 0.017 | 1.001 | 3055 | 3489 |
| ID (d_batch) | 0.145 | 0.017 | 0.115 | 0.182 | 1.000 | 4540 | 6374 |
| dyad (Intercept) | 0.019 | 0.002 | 0.015 | 0.023 | 1.002 | 2254 | 4976 |
| dyad (d_season) | 0.041 | 0.003 | 0.036 | 0.046 | 1.001 | 3907 | 6449 |
| dyad (d_date) | 0.006 | 0.003 | 0.000 | 0.012 | 1.007 | 872 | 2856 |
| dyad (d_time) | 0.006 | 0.002 | 0.001 | 0.010 | 1.006 | 1306 | 1690 |
| dyad (d_bodyregion) | 0.031 | 0.002 | 0.026 | 0.036 | 1.000 | 4125 | 6872 |
| dyad (d_s.distance) | 0.018 | 0.001 | 0.015 | 0.021 | 1.001 | 3782 | 7017 |
| dyad (d_s.volume) | 0.012 | 0.002 | 0.008 | 0.015 | 1.001 | 2756 | 3704 |
| dyad (d_subsamples) | 0.015 | 0.002 | 0.012 | 0.019 | 1.002 | 2898 | 4596 |
| dyad (d_batch) | 0.086 | 0.004 | 0.078 | 0.095 | 1.000 | 5176 | 8510 |

Table S6: Estimates for all fixed and random terms for the compositional model on social attributes. CrI = Credible Interval. ESS = effective sample size.

| **fixed term** | **Estimate** | **Est.Error** | **2.5% CrI** | **97.5% CrI** | **Rhat** | **bulk ESS** | **tail ESS** |
| --- | --- | --- | --- | --- | --- | --- | --- |
| Intercept | -0.003 | 0.164 | -0.326 | 0.321 | 1.002 | 2990 | 4034 |
| elo score | 0.000 | 0.009 | -0.018 | 0.018 | 1.001 | 10948 | 6198 |
| sex (male) | 0.000 | 0.022 | -0.042 | 0.043 | 1.000 | 9709 | 5772 |
| age | 0.000 | 0.011 | -0.021 | 0.021 | 1.000 | 10765 | 6682 |
| group (F) | 0.000 | 0.038 | -0.073 | 0.073 | 1.000 | 6257 | 5939 |
| group (H) | 0.001 | 0.038 | -0.074 | 0.075 | 1.000 | 5793 | 5660 |
| season (spring) | 0.002 | 0.114 | -0.225 | 0.225 | 1.000 | 3150 | 4068 |
| season (summer) | 0.001 | 0.113 | -0.223 | 0.224 | 1.001 | 3400 | 4811 |
| time | 0.000 | 0.009 | -0.018 | 0.018 | 1.001 | 11210 | 6215 |
| bodyregion (lower) | -0.001 | 0.094 | -0.182 | 0.182 | 1.000 | 5589 | 5075 |
| bodyregion (mixed) | 0.000 | 0.093 | -0.180 | 0.183 | 1.000 | 5615 | 5251 |
| bodyregion (upper) | -0.001 | 0.095 | -0.186 | 0.183 | 1.000 | 5692 | 5283 |
| s.distance | 0.000 | 0.023 | -0.045 | 0.044 | 1.001 | 2774 | 3951 |
| s.volume | 0.000 | 0.060 | -0.118 | 0.116 | 1.006 | 539 | 1202 |
| subsamples | 0.000 | 0.015 | -0.028 | 0.029 | 1.000 | 7401 | 6173 |
| **random term** | **Estimate** | **Est.Error** | **2.5% CrI** | **97.5% CrI** | **Rhat** | **bulk ESS** | **tail ESS** |
| sample (Intercept) | 0.007 | 0.005 | 0.000 | 0.019 | 1.001 | 5732 | 3816 |
| compound (Intercept) | 1.500 | 0.096 | 1.320 | 1.695 | 1.003 | 2217 | 3112 |
| compound (elo score) | 0.017 | 0.012 | 0.001 | 0.043 | 1.002 | 3304 | 3855 |
| compound (sex) | 0.102 | 0.019 | 0.063 | 0.138 | 1.001 | 2819 | 3082 |
| compound (age) | 0.052 | 0.017 | 0.012 | 0.083 | 1.002 | 1709 | 1218 |
| compound (group) | 0.290 | 0.016 | 0.261 | 0.321 | 1.001 | 3772 | 4889 |
| compound (season) | 1.060 | 0.042 | 0.982 | 1.146 | 1.002 | 2698 | 4371 |
| compound (time) | 0.040 | 0.018 | 0.004 | 0.072 | 1.003 | 1657 | 2624 |
| compound (bodyregion) | 0.133 | 0.018 | 0.099 | 0.167 | 1.001 | 2690 | 3865 |
| compound (s.distance) | 0.262 | 0.018 | 0.229 | 0.298 | 1.000 | 2989 | 5028 |
| compound (s.volume) | 0.810 | 0.042 | 0.734 | 0.899 | 1.002 | 1140 | 2388 |
| compound (subsamples) | 0.128 | 0.015 | 0.099 | 0.157 | 1.000 | 3441 | 5449 |
| ID (Intercept) | 0.007 | 0.005 | 0.000 | 0.020 | 1.000 | 5272 | 3378 |
| ID (season) | 0.007 | 0.005 | 0.000 | 0.020 | 1.001 | 6584 | 4186 |
| ID (time) | 0.007 | 0.005 | 0.000 | 0.019 | 1.001 | 6170 | 3882 |
| ID (bodyregion) | 0.007 | 0.005 | 0.000 | 0.019 | 1.001 | 5767 | 3616 |
| ID (s.distance) | 0.007 | 0.005 | 0.000 | 0.020 | 1.000 | 6069 | 3831 |
| ID (s.volume) | 0.007 | 0.005 | 0.000 | 0.020 | 1.000 | 5974 | 3933 |
| ID (subsamples) | 0.007 | 0.006 | 0.000 | 0.021 | 1.001 | 5094 | 3889 |
| date (Intercept) | 0.007 | 0.005 | 0.000 | 0.020 | 1.000 | 5750 | 3828 |
| date (elo score) | 0.007 | 0.005 | 0.000 | 0.019 | 1.000 | 6316 | 3602 |
| date (sex) | 0.007 | 0.005 | 0.000 | 0.019 | 1.000 | 5381 | 3579 |
| date (age) | 0.007 | 0.005 | 0.000 | 0.019 | 1.000 | 5632 | 3566 |
| date (group) | 0.007 | 0.005 | 0.000 | 0.019 | 1.000 | 5834 | 3434 |
| date (time) | 0.007 | 0.005 | 0.000 | 0.019 | 1.000 | 5882 | 3499 |
| date (bodyregion) | 0.007 | 0.005 | 0.000 | 0.019 | 1.000 | 5275 | 3463 |
| date (s.distance) | 0.007 | 0.005 | 0.000 | 0.019 | 1.001 | 5856 | 3897 |
| date (s.volume) | 0.007 | 0.005 | 0.000 | 0.020 | 1.000 | 6369 | 4406 |
| date (subsamples) | 0.007 | 0.005 | 0.000 | 0.020 | 1.000 | 5913 | 3819 |
| batch (Intercept) | 0.011 | 0.009 | 0.000 | 0.035 | 1.000 | 5531 | 4316 |
| tube (Intercept) | 0.007 | 0.005 | 0.000 | 0.019 | 1.000 | 5914 | 3912 |

Table S7: List of compounds potentially associated with sex and age differences in chemical profiles of Barbary macaque samples. Credible intervals of compounds listed in association with sex overlapped between the sexes, but the estimate (intercept) for one sex was outside the credible interval of the other. Similarly, credible intervals of compounds listed in association with age overlapped 0, but did so by < 20 % of the respective credible interval. RT = retention time (in min.), base peak describes the largest mass fragment of the peak. SI = similarity index (from 0 - 100) for the match between mass spectra and their respective best hit in the NIST library.

| **trait** | **RT** | **base peak** | **best NIST library hit** | **SI** | **substance class** | **found in** |
| --- | --- | --- | --- | --- | --- | --- |
| age | 4.44 | 59 | Formamide, N-methyl- | 95 | amide | humans ^8^ |
| age | 6.36 | 91 | Ethylbenzene | 85 | aromatic hydrocarbon | humans ^8–10^ |
| sex | 9.25 | 94 | Undecanal | 91 | aldehyde | humans ^8,11,12^, mandrills ^13^ |
| sex | 10.07 | 93 | 3-Carene | 92 | monoterpene | human excretions ^9^ |
| sex | 14.19 | 67 | unidentified | < 80 |  |  |
| age | 17.73 | 85 | unidentified | < 80 |  |  |
| sex | 30.84 | 119 | unidentified | < 80 |  |  |
| sex | 31.34 | 119 | unidentified aldehyde | < 80 | aldehyde |  |
| sex | 44.50 | 103 | Unidentified ester | < 80 | ester |  |

*Additional references:*

1.	Brauch, K. *et al.* Sex-specific reproductive behaviours and paternity in free-ranging Barbary macaques (*Macaca sylvanus*). *Behav. Ecol. Sociobiol.* **62**, 1453–1466 (2008).

2.	Nsubuga, A. M. *et al.* Factors affecting the amount of genomic DNA extracted from ape faeces and the identification of an improved sample storage method. *Mol. Ecol.* **13**, 2089–2094 (2004).

3.	Westphal, H. Paternity assessment in Barbary macaques. (Leipzig University, Leipzig, 2020).
